# Features Of Hippocampal Astrocytic Domains And Their Spatial Relation To Excitatory And Inhibitory Neurons

**DOI:** 10.1101/2020.05.25.114348

**Authors:** Ron Refaeli, Adi Doron, Aviya Benmelech-Chovav, Maya Groysman, Tirzah Kreisel, Yonatan Loewenstein, Inbal Goshen

## Abstract

The mounting evidence for the involvement of astrocytes in neuronal circuits function and behavior stands in stark contrast to the lack of detailed anatomical description of these cells and the neurons in their domains. To fill this void, we imaged >30,000 astrocytes in cleared hippocampi, and employed converging genetic, histological and computational tools to determine the elaborate structure, distribution and neuronal content of astrocytic domains. First, we characterized the spatial distribution of >19,000 astrocytes across CA1 lamina, and analyzed the detailed morphology of thousands of reconstructed domains. We then determined the excitatory content of CA1 astrocytes, averaging above 13 pyramidal neurons per domain and increasing towards CA1 midline. Finally, we discovered that somatostatin neurons are found in close proximity to astrocytes, compared to parvalbumin and VIP inhibitory neurons. This resource expands our understanding of fundamental hippocampal design principles, and provides the first quantitative foundation for neuron-astrocyte interactions in this region.

## INTRODUCTION

Recent years have seen a surge in development of new methods for neuroscience research, enabling detailed functional investigation of specific cell populations, alongside their elaborate anatomical characterizations. In astrocyte research, manipulations using chemogenetic and optogenetic tools were successfully integrated into the field, resulting in a new understanding of how these cells are an integral part of the circuits controlling emotional and cognitive behavior (Yu et al., 2020). This recent progress in deciphering the functional significance of astrocytes stands in stark contrast to the lack of comprehensive anatomical description of these cells, and of the neurons in their domains. Compared to the rich anatomical data available describing neurons, no large-scale characterization of the structure and spatial distribution of astrocytes has been performed.

Two decades ago astrocytes in the hippocampus were shown to exhibit a unique spatial organization: In contrast to the organization of neuronal dendrites, which are spatially intermingled, astrocytic processes display minimal overlap (Bushong et al., 2002). This finding was later replicated both in the hippocampus (Bushong et al., 2003, 2004; Livet et al., 2007; Ogata and Kosaka, 2002; Wilhelmsson et al., 2006; Xu et al., 2014) and in the cortex (Grosche et al., 2013; Halassa et al., 2007; Lopez-Hidalgo et al., 2016; Oberheim et al., 2008; Wilhelmsson et al., 2006) in different species, using various experimental techniques. The common staining procedures for astrocytes tag only their main processes, representing no more than 15% of their total volume (Bushong et al., 2002), and hence do not expose their true elaborate 3D morphology. To obtain a detailed description of astrocyte discrete domains, early works employed dye filling in CA1 astrocytes (e.g. (Bushong et al., 2002; Ogata and Kosaka, 2002; Wilhelmsson et al., 2006), and later ones induced the expression of fluorophores in astrocytes using transgenic animals (e.g. (Halassa et al., 2007; Livet et al., 2007) or viral vectors (Halassa et al., 2007; Jones et al., 2018). However, these studies were performed in brain slices 100-300μm thick, resulting in most astrocytes being truncated due to their large territories, thus radically restricting the number of analyzed cells. Indeed, the number of reconstructed astrocytes in all of the above studies ranged from less than twenty to a few hundred at most.

Recently developed brain clearing techniques allow large scale, three dimensional, single-cell resolution characterization of multiple cell types simultaneously (Gradinaru et al., 2018; Treweek and Gradinaru, 2016; Ye et al., 2016). Such methods can be adopted to investigate astrocytic domains, and indeed a few studies have already used transgenic mice to show that fluorescent astrocytes can be imaged in the cleared mouse cortex (Chai et al., 2017; Gaire et al., 2018; Lanjakornsiripan et al., 2018; Miller and Rothstein, 2016). However, the number of reconstructed astrocytes in these works remained very low, and their territories were defined in most cases by their envelope surfaces, rather than by the detailed morphology of their filaments. Thus, a thorough characterization of the morphology and distribution of astrocytic domains in the hippocampus is currently missing.

A crucial open question in the investigation of astrocytic domains is the content of different neuronal cell types within them. In principle, the average number of any specific neuronal cell type per domain can be fairly accurately calculated by simply dividing the total number of neurons by the total number of astrocytes, even in thin slices. However, such an average estimation relies on the incorrect assumption that astrocytes are similar in shape, volume, and distribution along grey matter lamina, and gives scant information regarding possible unique spatial interactions (e.g. proximity of astrocytes to specific cell types) and domain-specific constrains on neuronal occupancy. Even when full domains of fluorescently-tagged astrocytes were imaged in the past, the common use of sparse astrocytic labeling hindered the accurate evaluation of the neuronal content of their domains, as neurons that are only partially engulfed by one astrocyte may in fact belong to the domain of an adjacent non-fluorescent astrocyte (Chai et al., 2017; Halassa et al., 2007; Octeau et al., 2018). Currently, no data describing the distribution of specific neuronal cell-types in astrocytic domains, or even its mean value are available. Imaging fluorescently tagged cell populations in clear brains now allows a thorough characterization of the spatial interactions between astrocytes and a variety of neuronal cell types.

In this work, we employed converging genetic, histological, imaging and computational tools to determine the elaborate structure, distribution and neuronal content of hippocampal astrocytic domains. We imaged over 30,000 hippocampal astrocytes, orders of magnitudes more than ever reported, and provide a quantitative characterization of the spatial distribution of over 19,000 astrocytes in CA1 lamina, as well as detailed morphological analysis of thousands of reconstructed astrocytes (>6,700). We offer the first cell-type specific quantitative analysis of the neuronal occupancy of astrocytic domains by pyramidal excitatory neurons, and of their proximity to parvalbumin (PV), VIP and somatostatin (SST) inhibitory neurons. This resource provides a large-scale description of fundamental design principles in the hippocampus, relating the distributions and spatial locations of astrocytes, pyramidal neurons and inhibitory neurons. Such data defines the range of possibilities for the inter-domain effects of astrocytes on neuronal activity, allowing improved interpretation of experimental data and facilitating the design of future experiments.

## RESULTS

### Large Scale Automatic Detection of Astrocytic Domains in 3D Volumes

The pioneering studies characterizing astrocytic domains in the hippocampus employed 100-300μm-thick hippocampal slices, which restricted the number of fully reconstructed astrocytes (less than twenty to a few hundred at most), as many of them were truncated (Bushong et al., 2003, 2004; Bushong et al., 2002; Livet et al., 2007; Ogata and Kosaka, 2002; Wilhelmsson et al., 2006; Xu et al., 2014), thus imaging of much larger volumes is necessary. To rigorously characterize the structural and spatial characteristics of astrocytic domains, and their genetically-defined neuronal content, 2-3 orders of magnitude larger quantities of analyzed cells are necessary.

To achieve this demanding goal, we employed viral vectors to fluorescently tag astrocytes densely in WT or transgenic mice. Fluorophores (tdTomato or eGFP) were expressed under the GFAP promoter with high penetrance (>96.7%; Figure S1A), and almost complete specificity (>97%; Figure S1B). We then rendered thick brain slices (4-5mm) transparent by CLARITY, and acquired high resolution images of optically-defined cubes (ranging in size from 520×370×520μm to 635×635×1500μm) from each slice (Figure 1A-B). Single astrocytic territories were then reconstructed (Figure 1C-F, Figure S1C-E, Supplementary movie 1), and compactly represented for further analysis (Figure 1G). This analysis was performed for all astrocytes in the entire imaging cube (Figure 1H,I), for 55 cubes, providing quantitative data of over 30,000 astrocytes.

**Figure 1.**
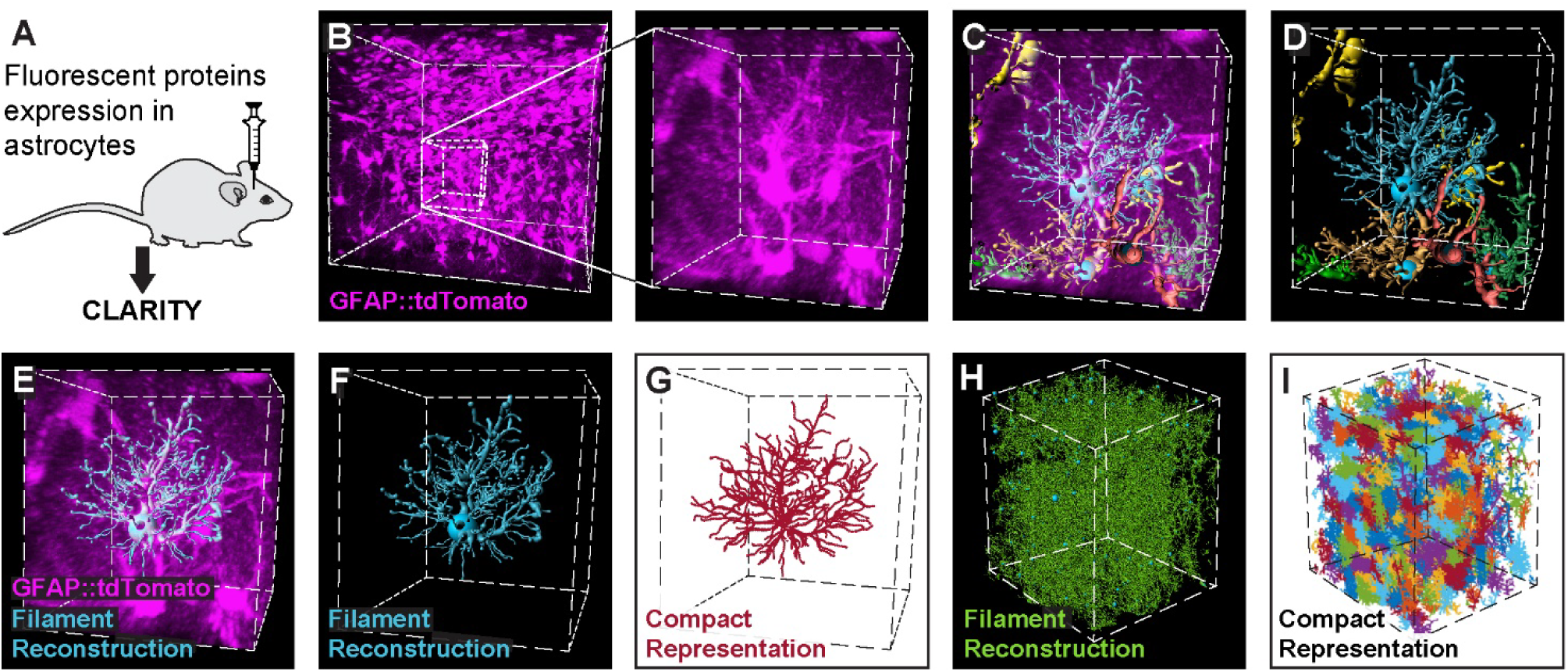
Automatic detection and reconstruction of astrocytic domains in large 3D volumes. (**A**) Wild type or transgenic mice were injected with viral vectors to induce the expression of fluorescent proteins in astrocytes, and thick hippocampal slices (4-5mm) were then made transparent by CLARITY. (**B**) Expression of tdTomato (purple) in hippocampal astrocytes is presented for a 450×450×450μm cube (left), and a zoomed-in 80×80×80μm cube excerpt from it (right). (**C-D**) Seven representative astrocyte filament reconstructions from this cube are shown (and see supplementary movie 1). Each astrocyte reconstruction, like the one shown in (**E-F**) was compactly represented (crimson) for further analysis (**G**). The process of filament reconstruction (green)(**H**) and compact representation (multicolor)(**I**) was performed on the entire imaging cube.

### Detailed Laminar Characterization of Astrocyte Domains in CA1

The large number of analyzed astrocytes obtained as detailed above was used to characterize the spatial distribution of astrocytes of different size and orientation along CA1 lamina with unprecedented detail. First, we quantified astrocytic density throughout CA1 lamina (Figure 2A-B). We found that the density of astrocytes in *stratum oriens* increases dramatically towards the pyramidal layer, whereas in the *stratum radiatum* the density is more uniform, with only a mild increase towards the pyramidal layer. In the pyramidal layer itself astrocytic density dramatically drops towards its middle (Figure 2C). The pyramidal layer varies in width, and hence we calculated the relative location of cells within it as the ratio between the distance to the lower edge and the total width. Then, we measured the size of the reconstructed astrocytes (Figure 2D). As depicted in Fig. 2E, the main finding is that starting ∼20µm from the pyramidal layer on both *stratum radiatum* and *stratum oriens*, astrocytic size decreases towards the middle of the pyramidal layer. Finally, we calculated the orientation of the same astrocytes relative to the surface of the CA1 pyramidal layer center (Figure 2F) and found it to be homogenous, on average, with all angles between 0º and 90º represented throughout hippocampal lamina (Figure 2G). This section provides a detailed analysis of the distribution of different astrocyte morphological characteristics that could potentially affect the neuronal content within these domains, which is what we sought to investigate next.

**Figure 2.**
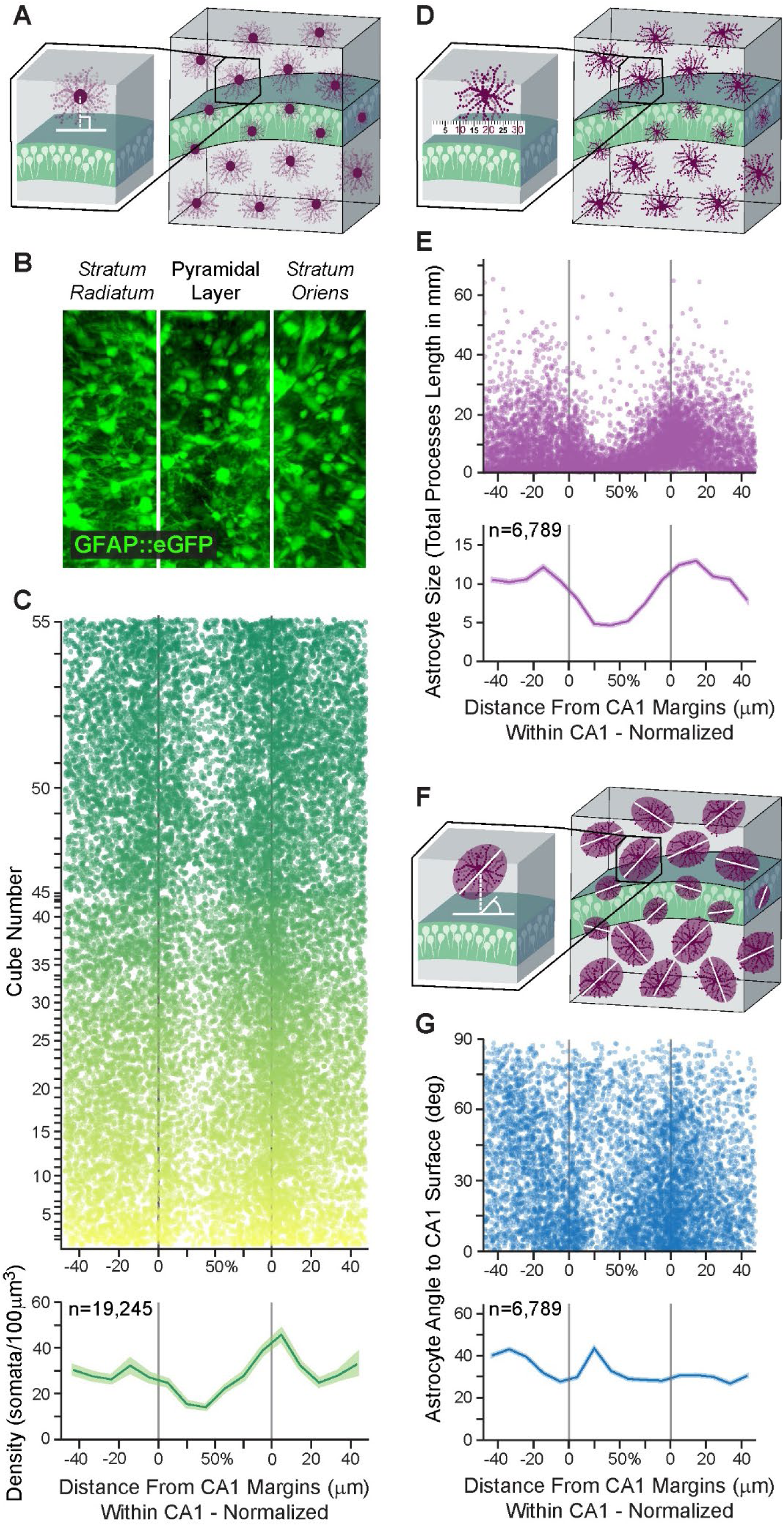
Large scale automatic detection of astrocytic domains and their detailed laminar characterization in CA1. (**A**) The distance of each astrocytic soma from the surface of the CA1 pyramidal layer was measured. (**B**) Representative z-projected cube showing astrocytes (green) in CA1. Lamina borders marked in white. (**C**) Astrocyte somata distribution (top; n=19,245 cells, each represented by a dot, from n=51 cubes, each displayed in a different semi-transparent color) and average astrocytic density along CA1 lamina (bottom). Density in both *stratum radiatum* and *stratum oriens* increases towards the pyramidal layer, and inside this layer density drops towards the middle from both sides. Average density (bottom) presented in bold red, with SEM shading. The distance within the CA1 pyramidal layer is normalized to correct for varying CA1 widths across samples. (**D**) Astrocyte size was calculated by measuring the total length of its processes. (**E**) The average size of hippocampal astrocytes (n=6,789 cells, each represented by a semi-transparent dot, from n=4 cubes) starts to drop at a ∼20µm distance from the pyramidal layer in both *stratum radiatum* and *stratum oriens*, and continues to decrease towards the middle. Average size (bottom) presented in bold purple, with SEM shading. (**F**) Astrocytic orientation was calculated relative to the surface of the CA1 pyramidal layer midline. (**G**) The orientation of the same astrocytes from panel D (each represented by a semi-transparent dot), relative to the surface of CA1 pyramidal layer. Average orientation (bottom) presented in bold blue, with SEM shading.

### Distribution of Excitatory Pyramidal Neurons in Astrocytic Domains

A large part of the research studying the functional interaction between astrocytes and pyramidal neurons was performed in CA1, and investigations into the role of astrocytes in memory also focused on this region (reviewed in (Santello et al., 2019)). However, almost nothing is known about the distribution of pyramidal neurons among the domains of the astrocytes around them. Given the variety of modulations of neuronal activity that can occur within each domain, and the proven spatial restriction of the effect of a single astrocyte manipulation to its domain’s vicinity (Henneberger et al., 2010), it is crucial to know how many pyramidal neurons are engulfed by each domain. To answer this question and quantify the spatial relations between astrocytes and pyramidal neurons, we co-labeled these populations as follows: Astrocytes were tagged in red using an AAV1-GFAP::tdTomato vector, and pyramidal neurons were labeled in green by a vector inducing the expression of eGFP in their nuclei (AAV5-CaMKII::H2B-eGFP)(Figure 3A-C, Figure S2A). This H2B nuclear tagging allows efficient separation of the pyramidal neurons, not possible when using a cytoplasm-filling fluorophore (Figure S2A-B). Astrocytic morphology was reconstructed as detailed above, and each neuronal soma was represented by a sphere (Figure 3D-E), and associated with a specific astrocytic domain (Figure 3F-H, Figure S2C; Supplementary movie 2. Supplementary stereoscopic movie 1). Neuronal content was determined only for astrocytes whose soma is in the pyramidal layer, or within 5µm from the edges of this layer. The number of pyramidal neurons per domain in these astrocytes ranged from 0 to 33 (with 50% between 10 and 17), averaging 13.7 (Figure 3I, Figure S2D-E). Only a weak positive correlation was found between the size of astrocytic domains and the number of neurons within them (F_(1,201)_=5.7, p<0.02, R^2^=0.027; Figure 3J). However, the number of neurons per domain was strongly correlated with the location of the astrocytes within CA1 (F_(1,201)_=76.6, p<0.001, R^2^=0.28), with increasing numbers of pyramidal neurons associated with astrocytic domains towards midline CA1 of the pyramidal cell layer (Figure 3K). Thus, when characterizing the number of pyramidal neurons in CA1 astrocytic domains, we found it to vary greatly (between 0 and 33), and be highly correlated with the location of the astrocyte relative to the pyramidal layer, and to a much lesser degree with the size of the astrocyte. As hippocampal activity is determined not only by pyramidal neurons, but is also continuously modulated by their neighboring inhibitory neurons, we next investigated the spatial interaction between astrocytes and the three most common inhibitory neurons in the CA1.

**Figure 3.**
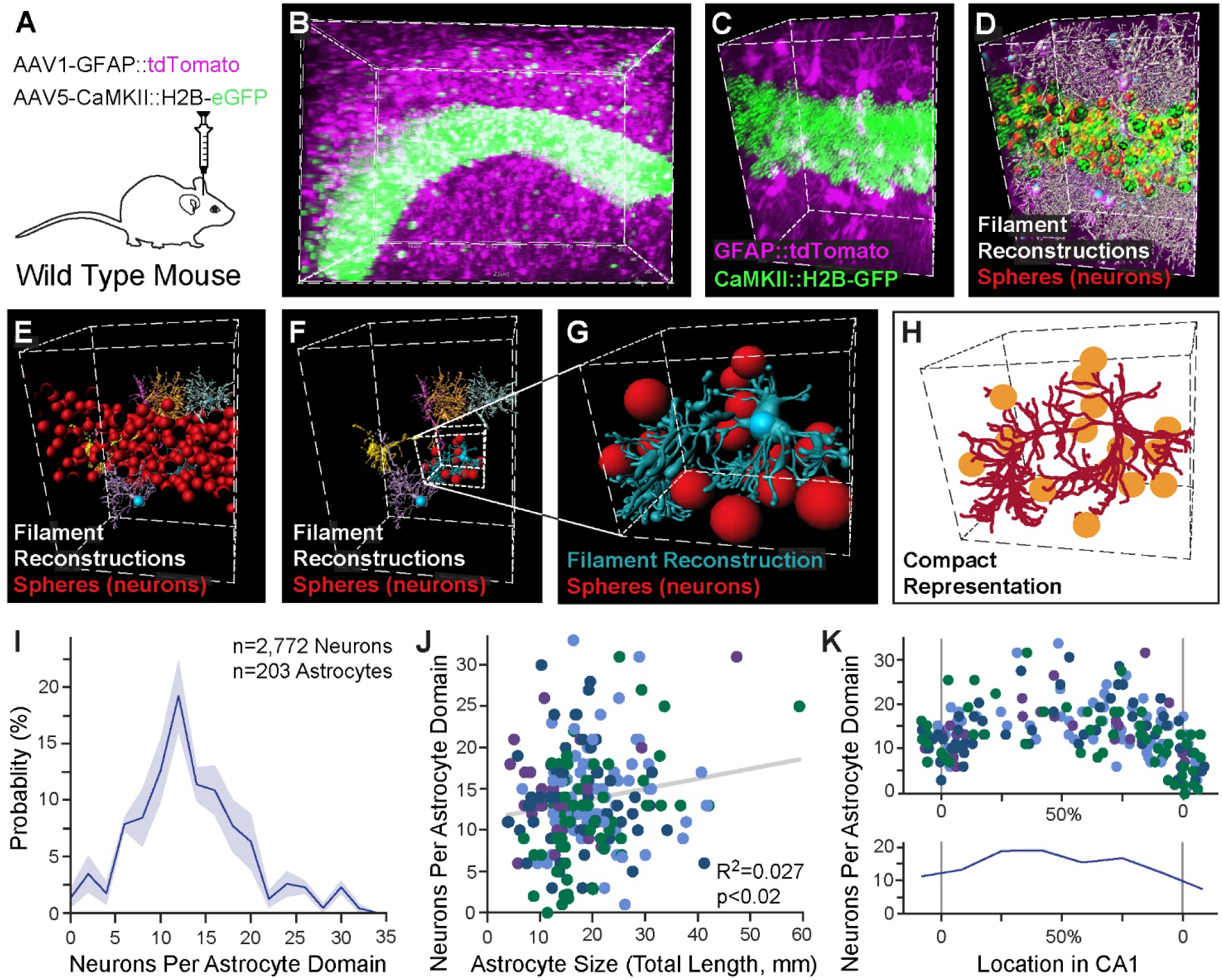
Automatic detection of excitatory neurons content and its distribution in astrocytic domains. (**A**) Mice were injected with viral vectors to induce the expression of the red fluorophore tdTomato in astrocytes, and the green fluorophore H2B-eGFP in pyramidal neurons nuclei, and thick slices (4-5mm) were then rendered transparent by CLARITY. Expression of tdTomato (purple) in hippocampal astrocytes and H2B-eGFP in pyramidal neurons (green) is presented for a 370×520×520μm cube (**B**), and a zoomed-in 80×100×250μm cube (**C**). (**D**) All astrocyte filament reconstructions from this cube are shown (white, and see supplementary movie 2), together with all neuronal somata (red). (**E**) Six representative filaments (blue, yellow, orange, pink, purple, grey) of astrocytes crossing the pyramidal layer are shown with all neuronal somata (red). (**F**) The same six filaments are shown only with the neuronal somata (red) associated with the blue astrocyte. (**G**) A zoomed-in image of the same reconstructed astrocyte and its associated neurons. (**H**) A compact matlab representation of the same astrocyte (crimson) and its associated neurons (yellow). (**I**) The binned distribution (bin size=2) of pyramidal neurons content of CA1 astrocytes (average=13.7; n=2,772 neurons in 203 astrocytic domains in 4 cubes). Average presented in bold blue, with SEM shading. (**J**) A significant albeit weak positive correlation is found between the size of the astrocyte and the number of pyramidal neurons in its domain (203 astrocytic domains, each represented by a dot, from n=4 cubes, each displayed in a different color). (**K**) The number of pyramidal neurons per astrocytic domain increases towards the middle of the pyramidal layer. Average number (bottom) presented in bold blue, with SEM shading.

### Spatial Associations Between Astrocytes and Inhibitory Neurons Subtypes

After characterizing the excitatory neuronal content of CA1 astrocytic domains, we sought to determine the spatial relations between astrocytes and inhibitory neurons in this region. The importance of astrocytes in GABAergic signaling in the normal and the diseased brain has been repeatedly demonstrated: Astrocytes respond to GABA with intracellular Ca^2+^ increase and can either mediate inhibition or convert GABAergic inhibition into glutamatergic excitation (Covelo and Araque, 2018; Perea et al., 2016). Furthermore, GABAergic cells depend on astrocytes for normal GABA synthesis (Robel and Sontheimer, 2016). A handful of studies directly investigated the functional interaction between astrocytes and inhibitory cell types, showing specificity in both astrocytic response to- and modulation of-distinct inhibitory populations: On the one hand, astrocytes show distinguishable responses to the activation of different inhibitory cell-types (Deemyad et al., 2018; Mariotti et al., 2018; Matos et al., 2018). For example, cortical astrocytes respond strongly to SST activation, but only weakly to PV activation (Mariotti et al., 2018). On the other hand, astrocytic manipulation can differentially modulate the effects of specific inhibitory neurons and not others (Deemyad et al., 2018; Matos et al., 2018; Perea et al., 2014; Tan et al., 2017). E.g., hippocampal astrocytes can detect SST activity and mediate its effects on pyramidal neurons, but they do not mediate the effect of PV cells in the same way (Matos et al., 2018). We found no reports on the interaction between VIP interneurons and astrocytes. While the functional interaction between astrocytes and neighboring inhibitory neurons has been investigated, the only available indirect quantification of spatial relationship between hippocampal astrocytes and neurons reports a sparse occupancy of neurons (of no defined type, but likely inhibitory due to their location) in the territories of *stratum radiatum* astrocytes (Chai et al., 2017). The genetic identity of these neurons is not known, but due to their location out of the pyramidal layer they are most likely inhibitory.

To gain insight into the spatial relations between CA1 astrocytes and inhibitory neurons, we co-labeled these populations as follows: Astrocytes were tagged in red, as before, using AAV1-GFAP::tdTomato. Inhibitory neurons were labeled by injecting a vector encoding a Cre-dependent green fluorophore (AAV5-EF1α-DIO-eGFP) to PV-Cre (Figure 4A), or VIP-Cre (Figure 4B), or SST-Cre (Figure 4C) transgenic mice (Hippenmeyer et al., 2005; Taniguchi et al., 2011). First, we set to quantify the spatial distribution of astrocytes and inhibitory neurons across CA1 lamina, and the interactions between them: In contrast to the distribution of astrocytic somata, which is highest around the margins of CA1 in all mouse strains (PV-Cre, VIP-Cre and SST-Cre; Figure 4D, as in Figure 2C), the three inhibitory cell types demonstrated clear differences in spatial distribution (Figure 4E, S3A). Quantifying these differences, we found that PV neurons were concentrated mainly in the pyramidal layer, with sparser expression in the *stratum oriens*. VIP cells were found almost exclusively in the pyramidal layer, especially close to its midline. SST cells presented a very different distribution, and were mostly located in the *stratum oriens* dendritic layer, with low expression in the pyramidal layer, and minimal expression in the *stratum radiatum* (Figure 4E, S3A). This spatial distribution is in line with previous reports (Atlas, 2007).

**Figure 4.**
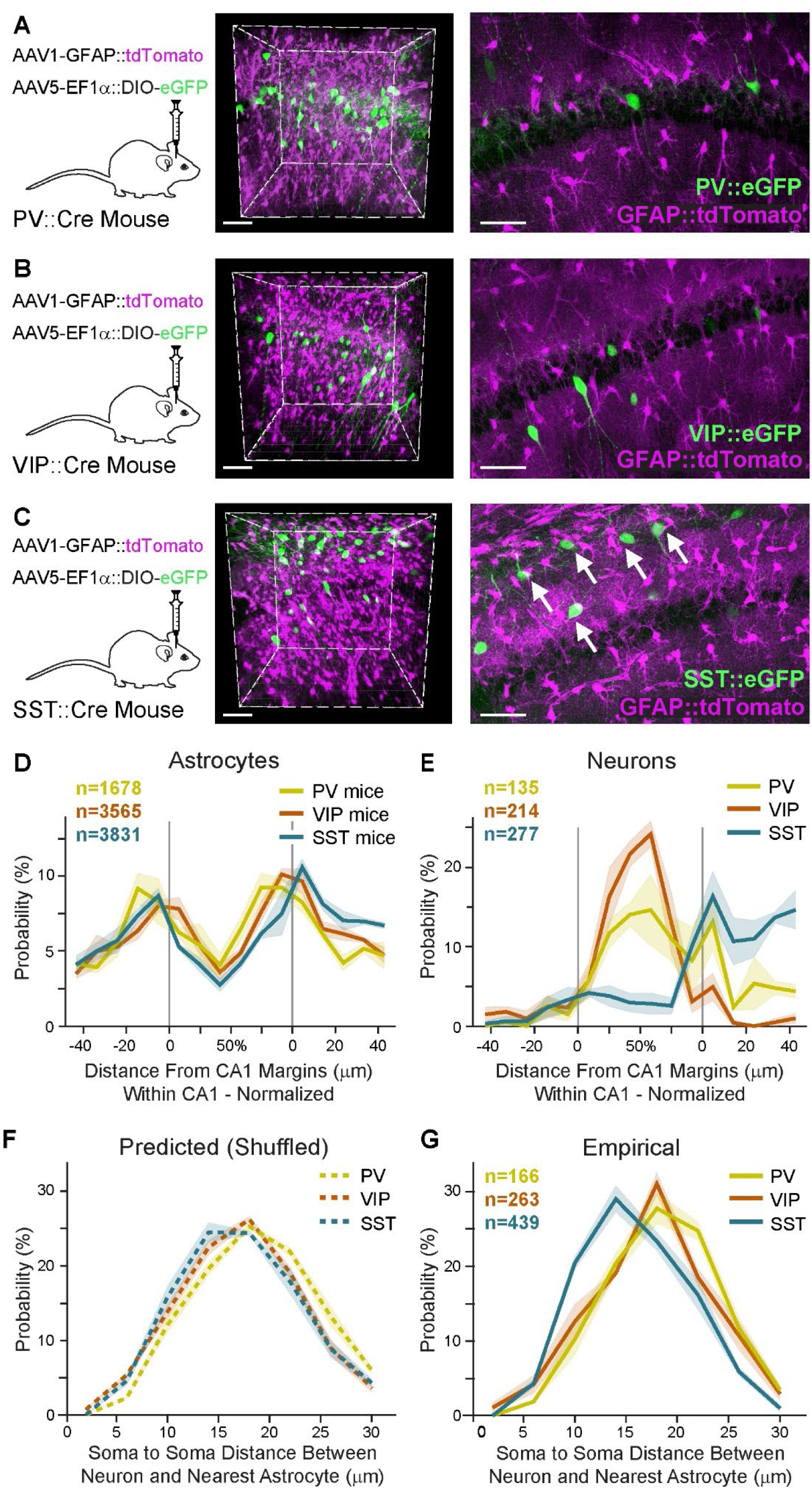
Unique proximity between astrocytes and somatostatin inhibitory neurons. AAV1-GFAP::tdTomato was injected into CA1 to tag astrocytes (purple), and AAV5-EF1α-DIO-eGFP to tag inhibitory neurons (green) in PV-Cre (n=3)(**A**), VIP-Cre (n=4)(**B**), or SST-Cre (n=4)(**C**) transgenic mice. Co-expression is shown in representative whole 400×400×450μm imaging cubes (middle), and one 25μm-thick optical plane from each cube (right). SST somata were observed in close proximity to astrocytes (white arrows in **C** *right*). (**D**) Astrocyte somata probability distributions in PV (yellow), VIP (orange) and SST (turquoise) mice, as a function of distance from the CA1 pyramidal layer surfaces, showing that astrocytes are highly concentrated on the borders of the pyramidal layer, and show low variance across the different samples. CA1 pyramidal layer borders are marked as in grey. (**E**) Spatial distribution of PV, VIP and SST somata, showing their mean density as a function of distance from the CA1 pyramidal layer margins. PV and VIP neurons are concentrated within the pyramidal layer, whereas SST neurons are mostly present in the dendritic layer in *stratum oriens*, with minor expression in the pyramidal layer. (**F**) The predicted (permutations-based) distribution of the distances between PV, VIP and SST somata to their nearest astrocyte soma (**G**). The empirical probability of SST somata to be in close proximity to the nearest astrocyte soma is significantly larger than that of PV and VIP (p<0.01; see Supplementary movie 3). Average density for each inhibitory cell type in bold color, with ±SEM shading. Scale bars: 70μm for cubes, 50μm for single planes.

The inhomogeneous distribution of astrocytes along CA1 lamina, together with the differences in the laminar distribution between the three inhibitory populations, may affect the spatial interactions between these cells. Indeed, when the expected distance between the inhibitory neurons somata and their nearest astrocyte soma was calculated for the three populations based on their spatial distribution, a significant difference was observed (F_(2,8)_=4.7, p<0.05), and post-hoc tests showed a significant difference between PV cells (p<0.05, with a higher expected soma-to-soma distance) and VIP and SST (Figure 4F). However, when we quantified the empirical distances between the soma center of each inhibitory cell to its nearest astrocyte soma center we found that SST neurons are located in much closer proximity to their neighboring astrocytes, relative to PV and VIP neurons (Figure 4G, 4C right, S3B-C; Supplementary movie 3). This striking difference was expressed by a significant difference between the average distance of the three populations to their closest astrocyte (F_(2,8)_=8.6, p<0.01), and post-hoc tests showed a significant difference between SST neurons and PV or VIP neurons (p<0.01 and p<0.05, respectively), but not between PV and VIP. Furthermore, when we compared the empirical distributions to the predicted ones, we found that the SST cells are, on average, closer to their neighboring astrocytes than anticipated (t_(3)_=2.9, p<0.05, paired t-test). No significant differences were found between empirical and predicted distributions for PV and VIP cells (p>0.05). The finding of a unique spatial correlation between astrocytes and SST inhibitory neurons corresponds with the reports of strong functional interaction between these cells.

## DISCUSSION

To fill the void between the mounting evidence for the functional significance of astrocytes in hippocampal function, and the lack of detailed anatomical characterization of these cells (Yu et al., 2020), we imaged over 30,000 hippocampal astrocytes, orders of magnitude more than ever reported, to provide a comprehensive quantitative characterization of their spatial distribution, detailed morphology, excitatory neurons content, and proximity to inhibitory neurons along CA1 lamina.

The investigation of astrocytic morphology to date was hindered mainly by technical barriers (Yu et al., 2020). In a recently published work, imaging of serially sectioned cortex using ChroMS microscopy followed the development of astrocytic clones and the 3D morphology of young and more mature astrocytes in these cortical clones (Clavreul et al., 2019). Brain clearing techniques now give access to large scale 3D imaging of fluorescently tagged cells, and specifically allow simultaneous imaging of more than one population.

The number of neurons per astrocytic domain varies greatly between regions, but even within each region estimating this number depends on the methodology. For example, in the striatum, the number of neurons was initially estimated to be around twenty (Chai et al., 2017), but using a different methodology this number was then reduced by half (Octeau et al., 2018). In both cases, sparse labeling of astrocytes was used, which can lead to over-estimation of the number of neurons per domain, as a neuron that is mostly engulfed by a nearby, non-fluorescing astrocyte may be erroneously associated with a fluorescent astrocyte. For that reason, our data was generated, by design, from samples with high-penetrance (>96%) expression of fluorophores in the astrocytic population, allowing a more accurate representation of the variance in neuronal content. We found that the number of pyramidal neurons somata is only mildly correlated with the size of the astrocyte, but is highly correlated to its location, i.e. astrocytes whose soma is closer to the pyramidal layer midline have more neurons in their domain. It could be hypothesized that this finding is caused by a bigger part of the surface area of these domains overlapping with the pyramidal layer, allowing more contact of the processes with neuronal somata.

When quantifying the spatial interaction between astrocytes and inhibitory neurons somata, we found that SST neurons are often ‘hugged’ by astrocytes, i.e. they are found in much closer proximity to neighboring astrocytes, compared to PV and VIP neurons. This unique interaction does not stem only from the fact that the density of both astrocytic and SST somata is high in the *stratum oriens*, but rather represents a unique design principle. Interestingly, this spatial proximity goes hand in hand with the reported unique functional interaction between these cells in the hippocampus, where astrocytes mediate the activity of SST (but not PV) cells (Matos et al., 2018).

The anatomical characterization of astrocytes was so far orthogonal to the investigation of their functional contribution to neuronal activity and behavior. However, the spatial distribution of these cells was shown to follow functional boundaries. For example, astrocyte processes are confined to barrel borders in the somatosensory cortex of mice, and to Cytochrome Oxidase (CO) blobs in human V1, and they show layer-specific molecular and morphological phenotype in somatosensory cortex (Eilam et al., 2016; Lanjakornsiripan et al., 2018). Thus, a functional significance for the local parcellation of the brain tissue by astrocytic territories exerting domain-specific effects on neuronal activity can be easily imagined. Indeed, LTP blockade in CA1 neurons, induced by calcium clamp in a single astrocyte was shown to be roughly restricted to its domain (Henneberger et al., 2010). We have recently shown that chemogenetic astrocytic activation in CA1 enhances memory and increases the number of active CA1 neurons recruited to support the acquisition of a new memory (Adamsky et al., 2018). Such an effect would require within-domain neuronal activity detection, and specific modulation of single neurons, but not others, within each domain.

To summarize, in this resource we performed a comprehensive quantitative characterization of the spatial distribution and morphology of CA1 astrocytes, in numbers that are orders of magnitudes larger than ever reported. We then provide the first cell-type specific quantitative analysis of the neuronal occupancy of astrocytic domains by pyramidal neurons, and of their proximity to three inhibitory neurons cell types. These results define the range of information that each astrocyte in the pyramidal layer is exposed to, and consequently the range of possibilities of within-domain modulation of single neurons. The functional significance of association with a specific astrocytic domain on spontaneous or evoked neuronal activity is yet to be demonstrated, and this resource provides the quantitative foundation upon which such future experiments can be designed.

## ACKNOWLEDGEMENTS

We thank the entire Goshen lab for their support. AD is supported by the Azrieli fellowship and the ELSC graduate students scholarship. This project has received funding from the European Research Council (ERC) under the European Union’s Horizon 2020 research and innovation programme (grant agreement No 803589), the Israel Science Foundation (ISF grant No. 1815/18), and the Canada-Israel grants (CIHR-ISF, grant No. 2591/18). We thank Ami Citri and Adi Kol for the critical reading of the manuscript.

## AUTHOR CONTRIBUTION

RR performed all CLARITY experiments, confocal and 2-photon imaging, and cell reconstructions. AD performed all data analysis. MG produced viral vectors. ABC and TK contributed to injections and histology. YL supervised data analysis. I.G. conceived and supervised all aspects of the project, and wrote the manuscript with input from all other authors.

## DECLARATION OF INTERESTS

The authors declare no competing interests.

## Supplementary Material

### METHODS

#### Mice

C57BL/6 WT mice, *Pv-IRES-Cre* (B6.129P2-Pvalb^tm1(cre)Arbr^/J – stock number 017320)(Hippenmeyer et al., 2005), Sst*-IRES*-Cre (Sst^tm2.1(cre)Zjh^/J - stock number 013044)(Taniguchi et al., 2011), and VIP*-IRES*-Cre (Vip^tm1(cre)Zjh^/J - stock number 010908)(Taniguchi et al., 2011), were used. 6-7 weeks old mice were group housed on a 12 hr light/dark cycle with *ad libitum* access to food and water. All mice were maintained under pathogen-free conditions in Tecniplast cages, on Teklad sani-chips (ENVIGO) bedding, at 20-24°C, and fed Teklad 2918SC (ENVIGO) pellets. Experimental protocols were approved by the Hebrew University Animal Care and Use Committee and met the guidelines of the National Institute of Health guide for the Care and Use of Laboratory Animals.

#### Stereotactic virus injection

Mice were anesthetized with isoflurane, and their head placed in a stereotactic apparatus (Kopf Instruments, USA). The skull was exposed and a small craniotomy was performed. To cover the entire dorsal CA1, mice were bilaterally microinjected using the following coordinates: Anteroposterior, −1.85mm from Bregma, mediolateral ± 1.4mm and dorsoventral −1.5mm. All microinjections were carried out using a 10µl syringe and a 34 gauge metal needle (WPI, Sarasota, USA). The injection volume and flow rate (0.1μl/min) were controlled by an injection pump (WPI). Following each injection, the needle was left in place for 10 additional minutes to allow for diffusion of the viral vector away from the needle track, and was then slowly withdrawn. The incision was closed using Vetbond tissue adhesive. For postoperative care, mice were subcutaneously injected with Tramadex (5mg/kg).

#### Viral vectors

AAV8-GFAP::eGFP, (UNC vector core). AAV1-GFAP::TdTomato, AAV5-EF1a-DIO-eYFP, AAV5-CaMKII-H2B-eGFP (ELSC Vector Core Facility, EVCF). Vectors were injected in a volume of 400 nl/site.

#### CLARITY

4-5mm thick hippocampal slices were cleared based on the protocol described by Ye et al (2016)(Ye et al., 2016). Briefly, mice were transcardially perfused with ice cold PBS followed by 4% PFA, brains were removed and kept in 4% PFA overnight at 4°C. Brains were then transferred to a hydrogel solution (PBS with: 2% acrylamide, bio-rad #161-0140; 0.1% Bisacrylamide, bio-rad #161-0142; 0.25% VA-044 initiator, Wako, 011-19365; 4% PFA) for 2 days. The samples were then degassed with N_2_ for 45 min and polymerized in 37°C for 3.5 hours. The samples were then washed overnight in 200mM NaOH-Boric buffer (sigma, #B7901) containing 8% SDS (sigma, #L3771), to remove PFA residuals. Samples were then stirred in a clearing solution (100mM Tris-Boric buffer, bio-lab, #002009239100 with 8% SDS) at 37°C for 3-4 weeks. After the samples became transparent, they were washed with PBST (PBS with 0.2% tritonX100; ChemCruz, #sc-29112A) for 24 hours at 37°C with mild shaking and for another 24 hours with fresh PBST 0.2% at RT. Finally, the samples were incubated in the refractive index matched solution Rapiclear (RI=1.47; SunJin lab, #RC147002) for 10 hours at 37°C and 2 days at room temperature before imaging.

#### Confocal Imaging

Cleared hippocampi were imaged by an Olympus scanning laser confocal microscope FV1000 using a water-immersion 20X NA 0.5 objective with 3.4mm working distance (UMPlanFL N). Confocal images were used for somata location identification and somata proximity analyses that did not require full morphological reconstruction.

#### 2-Photon Imaging

2-Photon imaging was performed using the Neurolabware 2-photon laser scanning microscope (Los Angeles, CA, USA). Excitation light from a Ti:sapphire laser (Chameleon Vision II, Coherent, and then Chameleon Discovery TPC, Coherent) scanned the sample using a 6215 galvometer and a CRS8 resonant mirror (Cambridge Technology). Emitted fluorescence light was detected by GaAsP photo-multiplier tubes (Hamamatsu, H10770-40) after bandpass filtering (Semrock). XYZ motion control was obtained using motorized linear stages, enabled via an electronic rotary encoder (KnobbyII). The Scanbox software, run on MATLAB, was used for microscope control and image acquisition. All images were acquired using a water immersion 16X objective (Nikon, 0.8 NA) with magnification of 2.8 or 3.4 to obtain 601×418µm or 516×366 µm fields of view, at 15.5 frames/second. In each sample, we acquired z-stacks with 0.937 µm intervals between planes, centered on the dorsal CA1 pyramidal layer. We scanned each plane 60 times and obtained its mean intensity projection image using FIJI. These images were concatenated into a single volumetric series for subsequent image analysis.

The chosen laser wavelength was optimized for astrocyte fluorescent emission, to enable filament reconstruction; Samples with GFP expressing astrocytes were scanned at 920 nm, which was adequate for detection of tdTomato expressing neuronal somas. Samples with tdTomato expressing astrocytes were scanned at 1040 nm, and a consecutive scan was conducted at 920 nm to enable neuronal detection in the green channel.

#### Image Analysis

##### Transforming size from CLARITY to non-clarified brains

To estimate the CLARITY induced expansion of the samples, we compared the size of 53 astrocyte somata in slices and in clarified brains.

We found CLARITY expansion factor be 1.89 in average, in all dimensions. This factor was used for all subsequent analyses to estimate the real-world distances and volumes.

##### Astrocyte filament reconstruction and compact representation

Astrocytes were segmented by constructing semi-automatic filaments in Imaris 9.1.2 (Bitplane, UK) while maintaining the same parameters for all samples (somata radius, dendrites radius, threshold, and maximal gap between points). Each astrocytic reconstruction was then extracted from Imaris to Matlab as 3D points coordinates with a connectivity matrix depicting the branching of the astrocyte. Detailed properties of all filaments such as length were extracted from Imaris as well.

##### Calculating the distance to CA1 pyramidal layer

Each cell of interest was represented as a single 3D point located at its soma center. Specifically, neuron somata were segmented using the spots option in Imaris, and then extracted as 3D points. Astrocyte somata locations were extracted from the filament reconstruction, defined as the filament starting point.

The CA1 pyramidal layer was masked using the manual surface option in Imaris, and then extracted to Matlab and smoothed using a moving mean. Gridded interpolants estimating the top and bottom surfaces of the CA1 pyramidal layer were created from the extracted boundaries, and each point of interest was evaluated accordingly to determine whether it resides below, above or inside this layer. Next, we found the nearest sampled point on the CA1 pyramidal layer boundary to the point of interest, and created a triangulated surface around it using its 4 neighboring grid points (above, below, right and left). Finally, the closest point on the triangulated surface to the point of interest was extracted, as well as the distance between them, using the point2trimesh function.

Because the CA1 pyramidal layer varies in size, we used normalization to present the distances of cells residing within it. First, we calculated the mean size of the CA1 pyramidal layer across all samples. Then, we measured the distances of each cell in this lamina to its boundaries, and calculated the normalized relative cell location as follows:

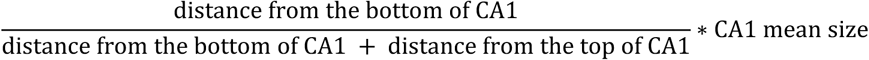

##### Determining astrocyte size

Astrocyte size was determined using the Imaris statistic “Filament Length”, which returns the sum of the lengths of all edges within the filament.

##### Calculating astrocyte orientation

To determine the orientation of each astrocyte, we calculated the first principle component based on the 3D point clouds that represent it. Next, we found the astrocyte centroid, and its nearest point on the midline surface of the CA1 pyramidal layer (obtained by averaging the bottom and top surfaces), as described above. We then calculated the angle between the normal to the CA1 surface and the principle component, and subtracted it from 90 degrees.

##### Allocating pyramidal neurons to astrocytic domains

To determine the pyramidal neurons occupancy of each astrocyte, we calculated the minimal distance between each neuron to the end points of all the astrocytes. The neuron was assigned to the astrocyte which had the closest end point relative to the neuron soma. To reduce the probability of including truncated domains in the analysis, we omitted astrocytes with somata that were close to the scanned volume edges (less than 20 µm from the edges) and/or far from the pyramidal layer (more than 5 µm from the CA1 surfaces). Neurons that were close to the scanned volume edges (less than 5 µm from the edges) and/or were assigned to an omitted astrocyte were also removed from the analysis.

##### Analyzing somata proximity

In each sample that contained tagged inhibitory neurons, we calculated the distance between each neuron soma to all the astrocyte somata. The minimal distance from each neuron soma to its nearest astrocyte somata in each cubic volume was measured, and used to calculate probability histograms.

#### Statistical analysis

Sample number (n) indicates the number of cells or imaging cubes in each experiment and is specified in the figure legends. Results were analyzed by one-way ANOVA, followed by LSD post-hoc tests, paired t-test, or a linear regression test, as applicable. All the statistical details of experiments can be found in the result section. Differences in means were considered statistically significant at p < 0.05. Analyses were performed using the IBM SPSS Statistics software (version 25).

##### Permutation tests

Permutation tests were conducted to test whether the empirical distributions are significantly different from predicted shuffled distributions. Specifically, we kept the astrocytes in their real locations, and simulated random neural locations. We permuted the neurons on the circumference of a 55.56 um circle, parallel to the CA1 pyramidal layer midline surface, such that each simulated neuron maintained its distance from this layer. This method enabled us to preserve the cellular spatial distributions relative to CA1, which are unique to each inhibitory cell type (see Figure 4). The simulation was run 1500 times, and the minimal distance between the permuted neuronal somata to astrocyte somata was calculated and binned similarly to the real data. When the simulated distances were smaller than the real minimal distance between the specific inhibitory cell to its nearest astrocyte soma, they were omitted, to avoid non-biologically plausible structures, i.e. cells with overlapping somas.

## SUPPLEMENTARY FIGURES

**Figure S1.**
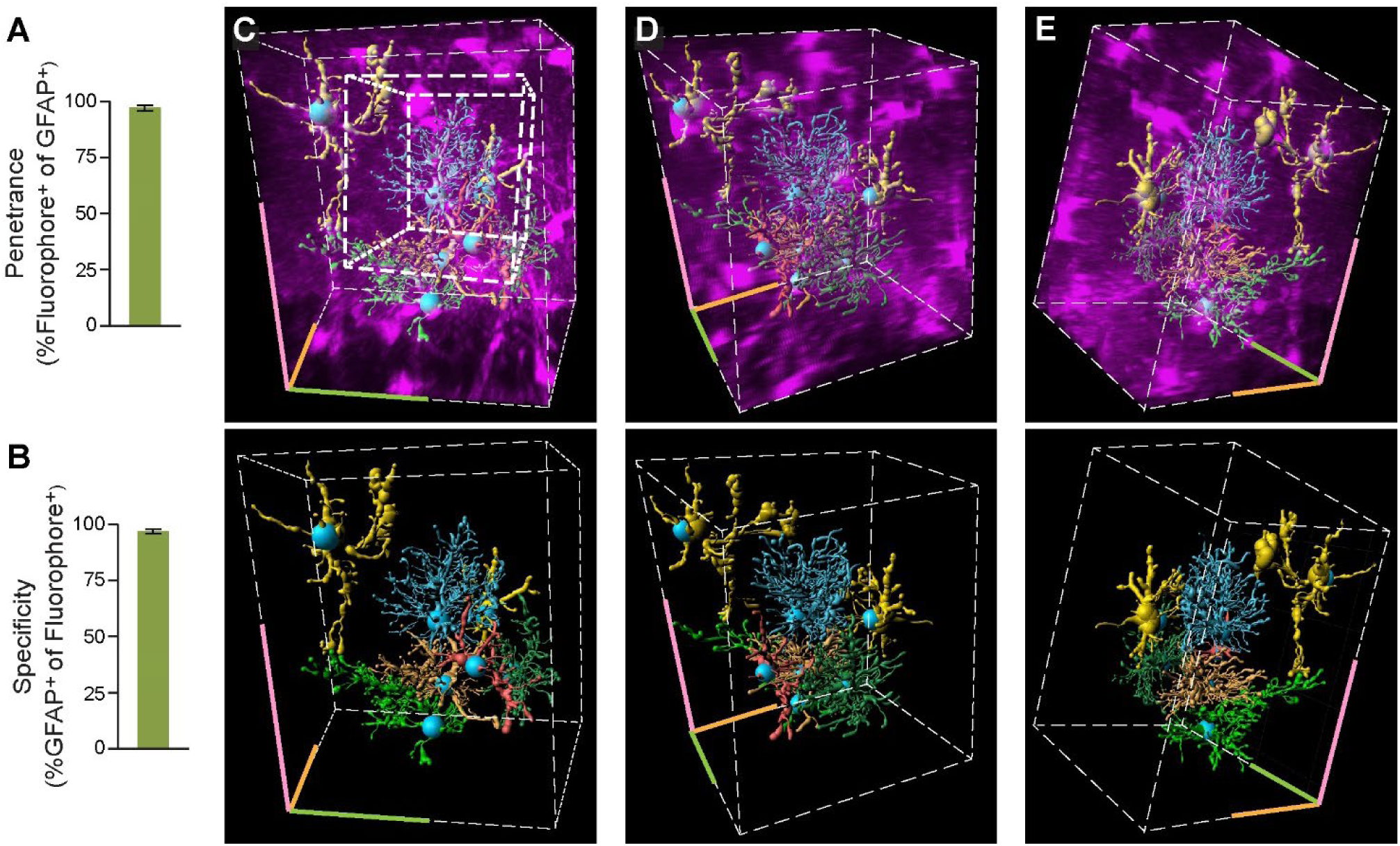
Representative 3D reconstruction of astrocytic domains. The fluorophores tdTomato and eGFP were expressed in >96.7% of CA1 astrocytes (292/301 of the GFAP cells expressed the fluorophore)(**A**), with >97% specificity (292/300 fluorescent cells were also GFAP positive)(**B**). (**C-E**) A zoomed-in part of the cube from Figure 1B, presenting the full structure of the seven astrocytes partially presented in Figure 1C,D (original cube from Figure 1C,D is marked in white on C top) from three different angles. To denote orientation, the same corner of the cube is marked in all images (pink-yellow-green). Note the elaborate structure and non-overlapping domains of these astrocytes.

**Figure S2.**
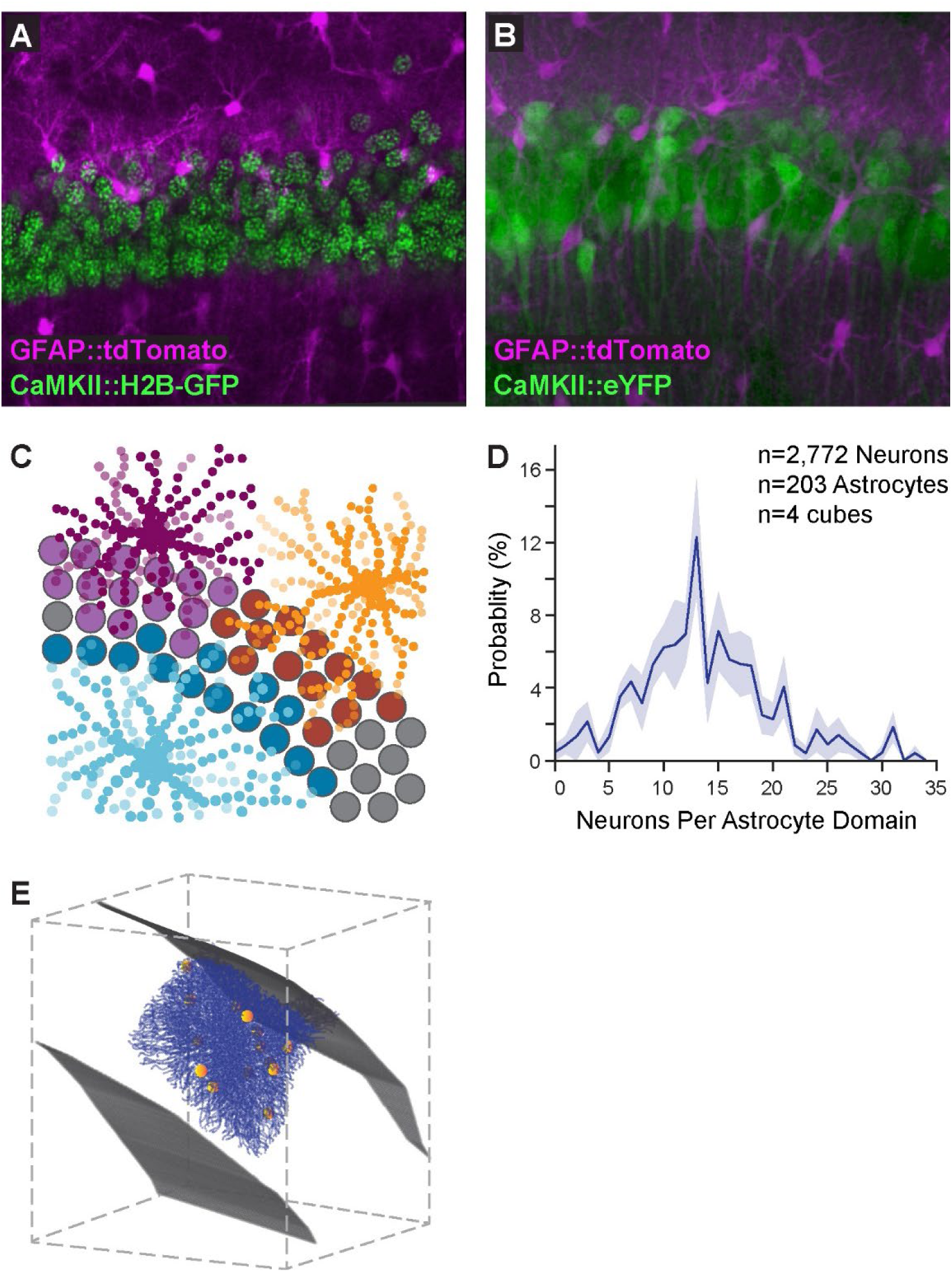
Distribution of excitatory neurons in astrocytic domains. (**A**) Expression of tdTomato (purple) in hippocampal astrocytes and H2B-eGFP in the nuclei of pyramidal neurons (green) is presented in a single plane 2D image. (**B**) When a cytoplasm-filling fluorophore (eYFP; green) was used under the same promoter, single neurons could not be automatically separated. (**C**) Illustration of neuronal allocation to nearest astrocyte. (**D**) A non-binned distribution of pyramidal neurons content of CA1 astrocytes (n=2,772 neurons in 203 astrocytic domains). Average presented in bold blue, with SEM shading. (**E**) An example astrocyte with its associated neurons: The soma of this astrocyte (processes in blue) is within the CA1 pyramidal layer (upper and lower pyramidal layer boundaries marked in grey), and 14 pyramidal neurons (somas marked in yellow) reside within its domain.

**Figure S3.**
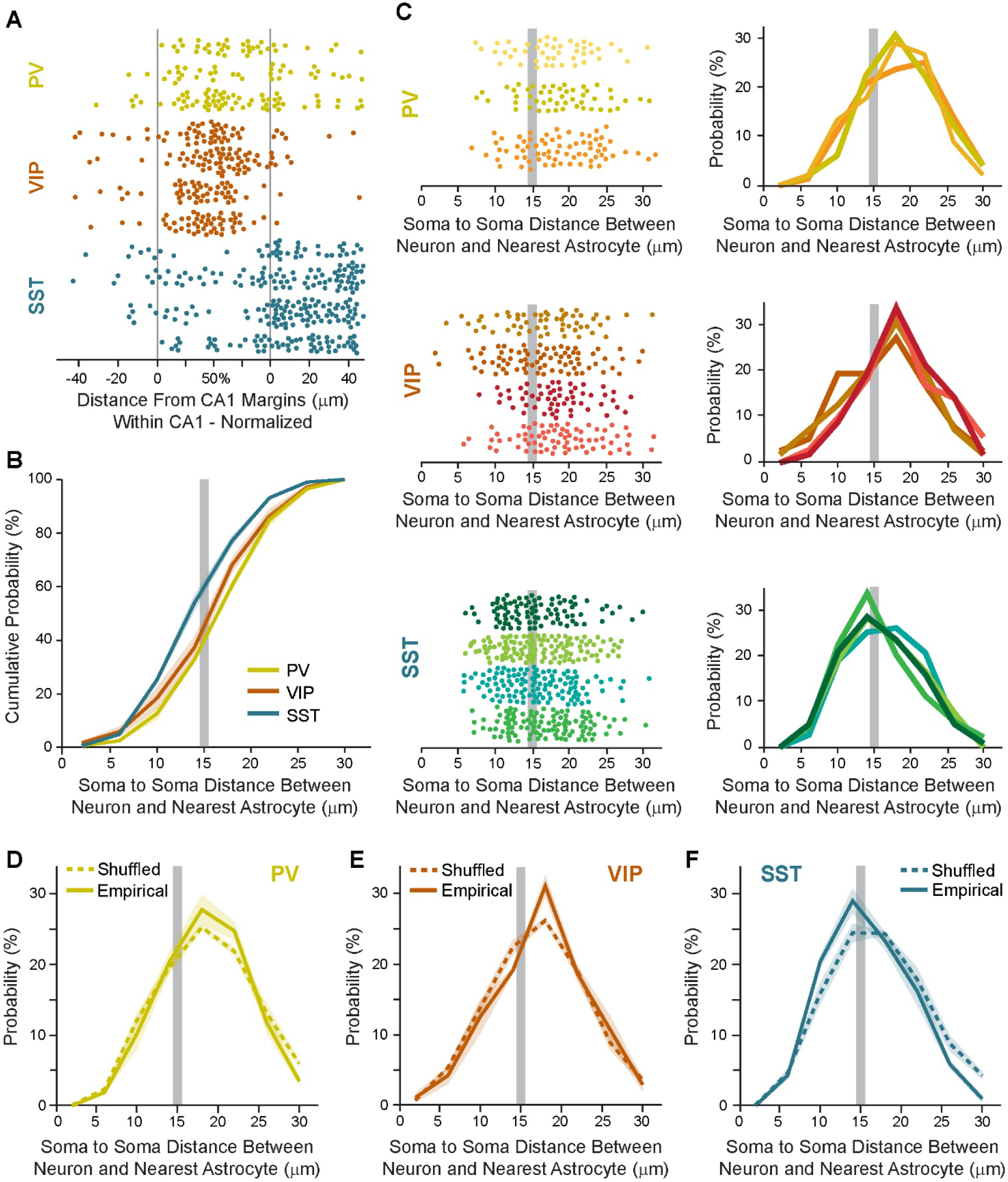
Low variance in the spatial associations between astrocytes and inhibitory neuron subtypes. (**A**) The raw data used to generate Figure 4E, showing the single cell spatial distribution in each sample, for PV (n=135), VIP (n=214) and SST (n=277) somata, as a function of distance from CA1 borders (grey lines). The distance within the CA1 pyramidal layer is normalized to correct for varying CA1 widths across imaging cube samples. (**B**) The cumulative distributions showing a higher probability of SST (turquoise) somata to be in close proximity to the nearest astrocyte soma compared to PV (yellow) and VIP (orange) neurons. (**C**) The data used to generate Figure 4G and panel B here, showing single somata (left) or distribution per imaging cube (right), for PV, VIP, and SST imaging cubes. (**D-F**) The empirical distance to nearest astrocyte distribution, presented together with the shuffled data distribution generated for PV, VIP and SST neurons (same as Figure 4F-G). Grey lines mark a 15μm soma-to-soma distance. Data presented as mean (bold) ±SEM (shadow).

## SUPPLEMENTARY MOVIES

### 2D MOVIES

**Movie 1. Automatic reconstruction of astrocytic domains**. The processes of hundreds of eGFP-expressing astrocytes (green, shown in the first 10 seconds, moving towards the depth of the cube) in a 3D cube from dorsal CA1 were reliably reconstructed into filaments (also green) from each cell core (blue, shown in the last 6 seconds, moving towards the periphery of the cube).

**Movie 2. Excitatory neurons content in astrocytic domains**. A full 520×370×520μm CA1 cleared cube, showing the nuclei of CA1 pyramidal cells (expressing H2B-eGFP, green) and astrocytes (expressing tdTomato, red) is shown, followed by plane-by-plane presentation, in which the filaments of a single astrocyte are reconstructed in blue. This single astrocyte is then presented in 3D, first together with all 1800 nuclei spots in CA1 (grey), and then only with the 19 somas associated with its domain.

**Movie 3. Proximity between astrocytes and somatostatin inhibitory neurons**. Astrocytes expressing tdTomato (magenta) and somatostatin (SST) inhibitory neurons expressing eGFP (green) in a 635×500×1300μm cleared CA1 cube. When entering the cube, the somata of five SST neurons and their neighboring astrocytes are marked by spots (same colors), to show the exceptional proximity between the two populations. Note that some SST somata are ‘hugged’ by more than one astrocyte.

### STEREOSCOPIC MOVIE

**Stereoscopic Movie 1. Excitatory neurons content in astrocytic domains**. A full 520×370×520μm CA1 cleared cube, showing astrocytes (expressing tdTomato, red) and the nuclei of CA1 pyramidal cells (expressing H2B-eGFP, green) and astrocytes (expressing tdTomato, red) is presented. *This movie is supported by all computer-based virtual reality system software*.

**Link to All Movies**: https://www.goshenlab.com/movies-1?pgid=kamhpf2e-f400a205-7208-4b84-a02b-27c7575075b8

## Notes

### Competing Interest Statement

The authors have declared no competing interest.

